## Supplemental Figures, Movie Captions for "A mechanism with severing near barbed ends and annealing explains structure and dynamics of dendritic actin networks"

**Movie S1. Simulated keratocyte lamellipodium.** Movie of simulated keratocyte lamellipodium in the rest frame of the cell (0-4 $\mu$m with the leading edge located at the top) for the enhanced end severing parameter set (Fig. 4). Oligomer fragments not shown. Fragments of filaments that appear correspond to annealing events and fragments that disappear to the creation of oligomers. End severing event are shown by disappearance of fragments near barbed ends. Uniform severing events can be identified by the appearance of pairs of pointed and barbed ends. Some filaments overlap one another as we do not have excluded volume interactions. Few filaments near the leading edge can be seen polymerizing after uncapping. These barbed ends were annealed to by a polymerizing oligomer created by uniform severing. The barbed end state of the oligomer was transferred to the filament resulting in a polymerizing filament away from the leading edge. Gray lines: actin filaments; red: Arp2/3 complex; yellow:free barbed ends; orange: capped barbed ends; blue: free pointed ends. Each frame is 0.1 s. Leading edge is 2 $\mu$ m wide.

**Movie S2. Simulated XTC lamellipodium.** Same as Movie S1 but using the parameters for XTC cells, enhanced end severing parameter set (Fig. 5).

**Movie S3. Simulated keratocyte actin SiMS.** Simulated actin SiMS for the enhanced end severing keratocyte parameter set (Fig. 4) with 0.01% of actin monomers tracked. Speckles are positioned based on the actin monomer location. Each frame is a collection of the appearance, disappearance, and motion within 1 s. Speckles that appeared within 1 s are colored in orange and located at their appearance location. Blue speckles remained associated to the network throughout the time range and are relocating with retrograde flow. Speckles that disappeared within this time frame are colored green and located at the disappearance location. Time stamp indicates the beginning of the 1 s interval.

**Movie S4. Simulated XTC cell actin SiMS.** Same as Movie S3 but for the enhanced end severing XTC parameter set (Fig. 5) with 0.05% of actin monomers tracked.

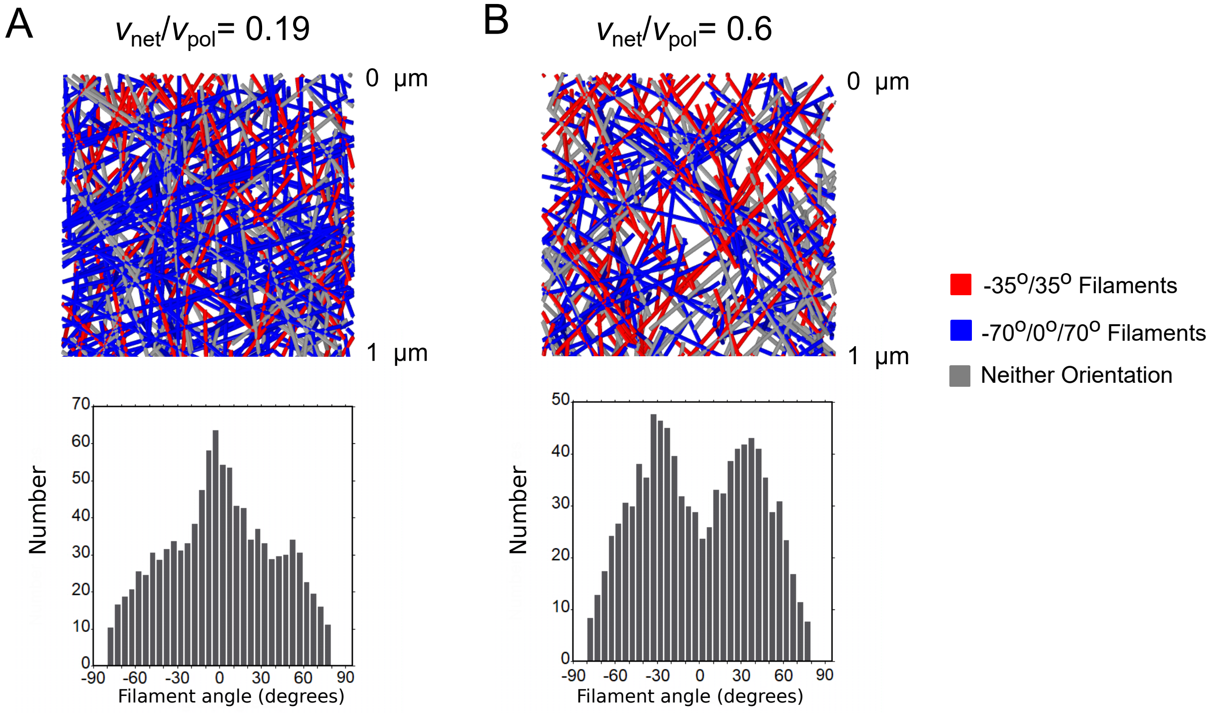


Figure S1: Addition of kinking does not effect the orientation pattern. (A) Top: 3D simulation snapshots, colored by orientation pattern, for $v_{net}/v_{pol}$ = 0.19 with kinking. Bottom: orientation distribution of filaments with a portion located within 1 $\mu$m of the leading edge, average of 5 simulations reaching steady state. Addition of kinking does not significantly alter the orientation distribution compared to the uniform 3D branching of Fig. 2C. (B) Same as panel A, for $v_{net}/v_{pol}$ = 0.6 with kinking, showing that kinking does not significantly change the orientation distribution compared to the uniform 3D branching of Fig. 2D. Snapshots are taken from simulations with a branching rate of 75 /sub/s and other parameters as in Table 1 (keratocyte parameters) with $k_{unif}^{sev} = 10^{-5}$ sub/s, and no annealing or enhanced end severing, $k_{anneal} = k_{end}^{sev} = l_{max}^{olig} = 0$.

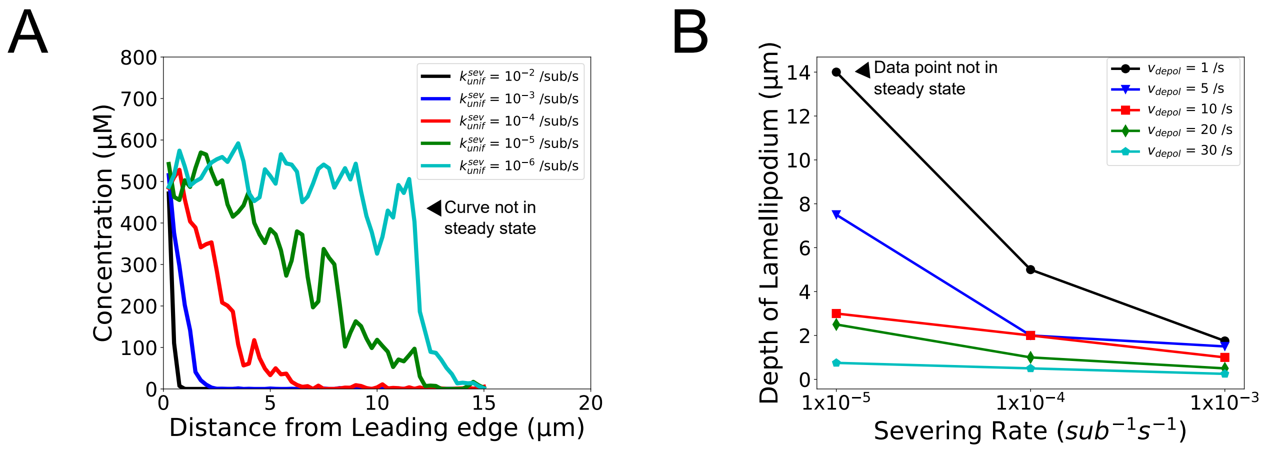


Figure S2: Simulated lamellipodium concentration and depth as a function of uniform severing rate $k_{unif}^{sev}$ and pointed end depolymerization rate $v_{depol}$. (A) F-actin concentration profile as function of $k_{unif}^{sev}$. (B) Lamellipodium depth as a function of $k_{unif}^{sev}$ and $v_{depol}$. The depth is the distance from the leading edge at which the concentration decreases by 50%. Both panels show the result of a single simulation at 300 s using XTC parameters listed in Table 1 (XTC parameters) but without annealing, enhanced end severing or uncapping, $k_{anneal} = k_{end}^{sev} = l_{max}^{olig} = k_{uncap} = 0$.

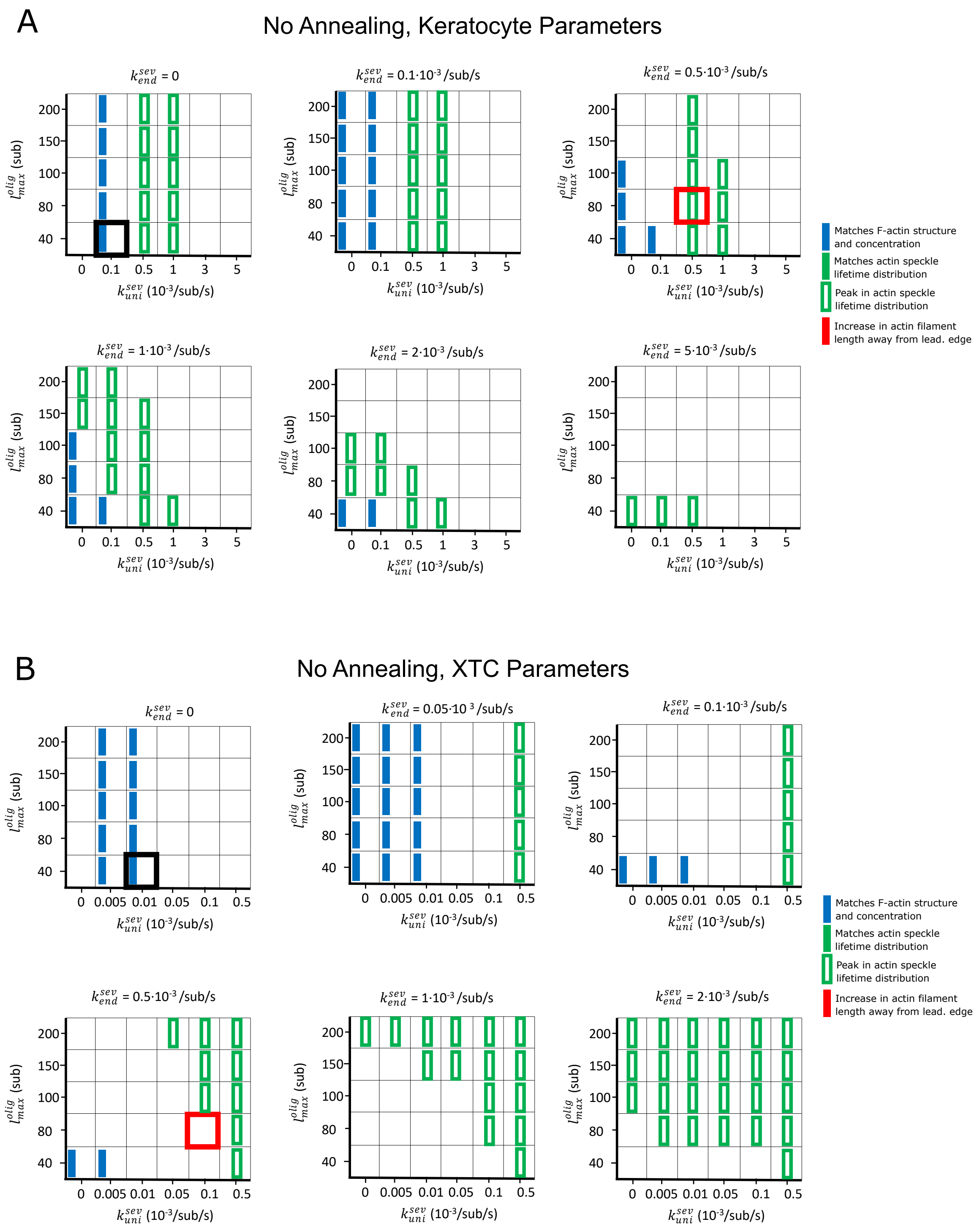


Figure S3: Parameter scan of uniform severing and enhanced end severing rates ($k^{sev}_{unif}$, $k^{sev}_{end}$) and maximum oligomer length ($l^{olig}_{max}$) without annealing. Parameters are listed in Table 1, without annealing or debranching: $k_{anneal} = k_{debr} = 0$. Boxes highlighted with thick black and red frames correspond to the parameters used in the black and red curves of Fig. 3. Blue rectangles indicate that F-actin, barbed end, and branch concentration profiles agree with experimental observations. Open green rectangles indicate that the simulated actin speckle lifetimes peaked at short times, but not within 2 s (keratocyte) or 4 s (XTC). No parameter combinations matched the SiMS speckle lifetime distribution and appearance or disappearance profiles (solid green) or provided a length increase away from the leading edge (red). (A) Keratocyte parameter scan. (B) Same as panel A, for XTC cells.

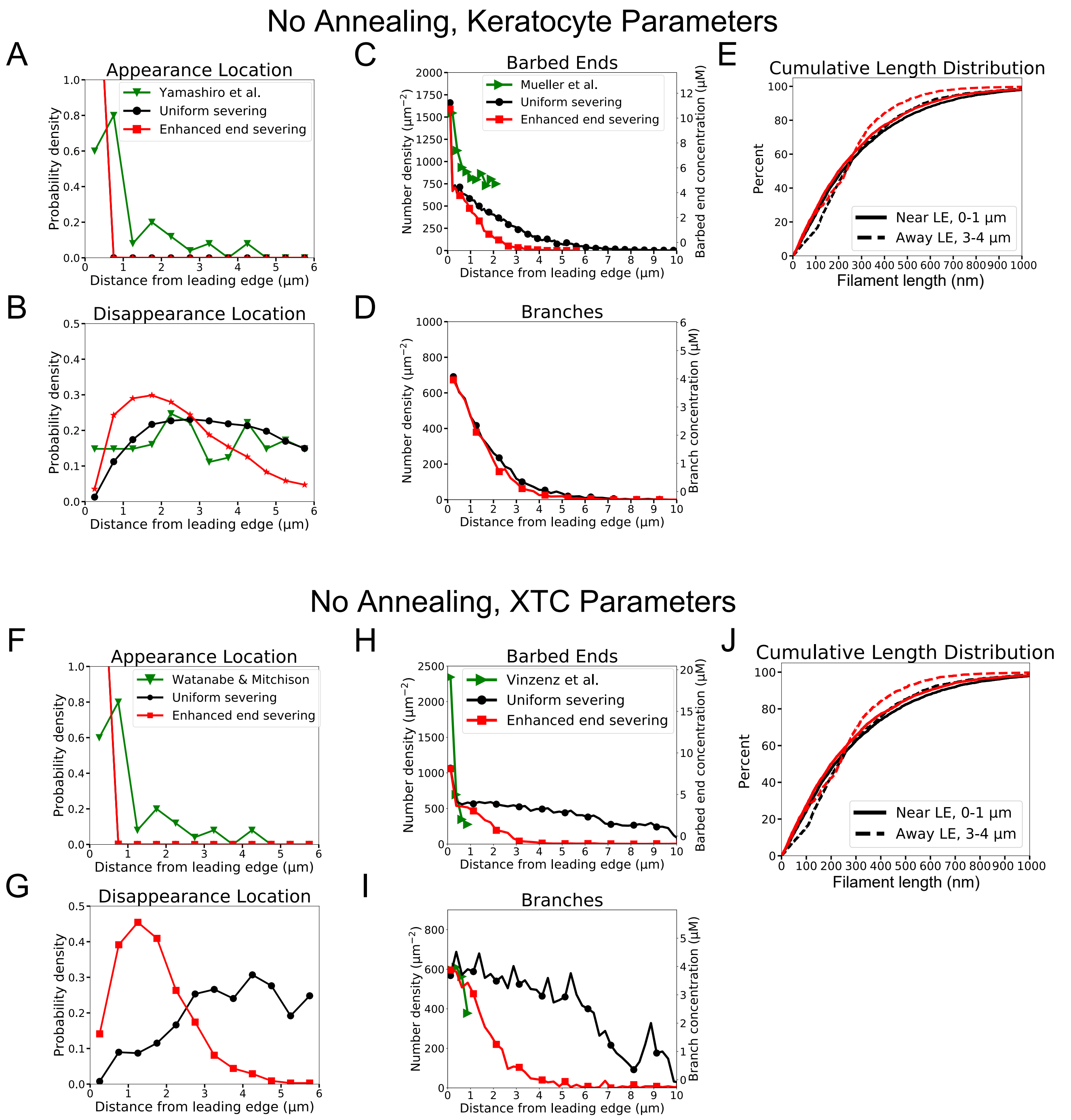


Figure S4: Quantification of model without annealing that did not reproduce the actin speckle lifetime and F-actin concentration profiles (continuation of Fig. 3). Results for models with uniform severing (black) or model with severing, enhanced near barbed end (red). (A-E) Keratocyte parameters as in Table 1 with $k^{sev}_{unif} = 10^{-4}$ /sub/s; $l^{olig}_{max} = 40$ sub (black) and $k^{sev}_{unif} = 5 \cdot 10^{-4}$ /sub/s; $k^{sev}_{end} = 5 \cdot 10^{-4}$ /sub/s; $l^{olig}_{max} = 80$ sub (red). (A) Simulated actin speckle appearance location and comparison to Yamashiro et al. (2018). Distributions were normalized within the indicated range, considering speckles with lifetimes longer than 2 s. (B) Same as C, for disappearance location. (C) Distribution of barbed ends and comparison to measurements in Mueller et al. (2017). (D) Concentration of simulated Arp2/3 complex branches. (E) Cumulative filament length distributions near ($0 - 1\mu m$, solid) and away from the leading edge ($3 - 4\mu m$, dashed). (F-J) Same as A-E for XTC cells and comparing to experimental data by Watanabe and Mitchison (2002) or Vinzenz et al. (2012). XTC parameters as in Table 1 with $k^{sev}_{unif} = 10^{-5}$ /sub/s; $l^{olig}_{max} = 40$ sub (black) or $k^{sev}_{unif} = 5 \cdot 10^{-4}$ /sub/s; $k^{sev}_{end} = 10^{-4}$ /sub/s; $l^{olig}_{max} = 80$ sub (red). (F,G) Distributions were normalized within the indicated range, considering speckles with lifetimes longer than 4 s. Data averaged over 5 independent simulations. Speckle data measured for speckles within 12 $\mu$m of the leading edge over a 20 s interval in steady state for each simulations.

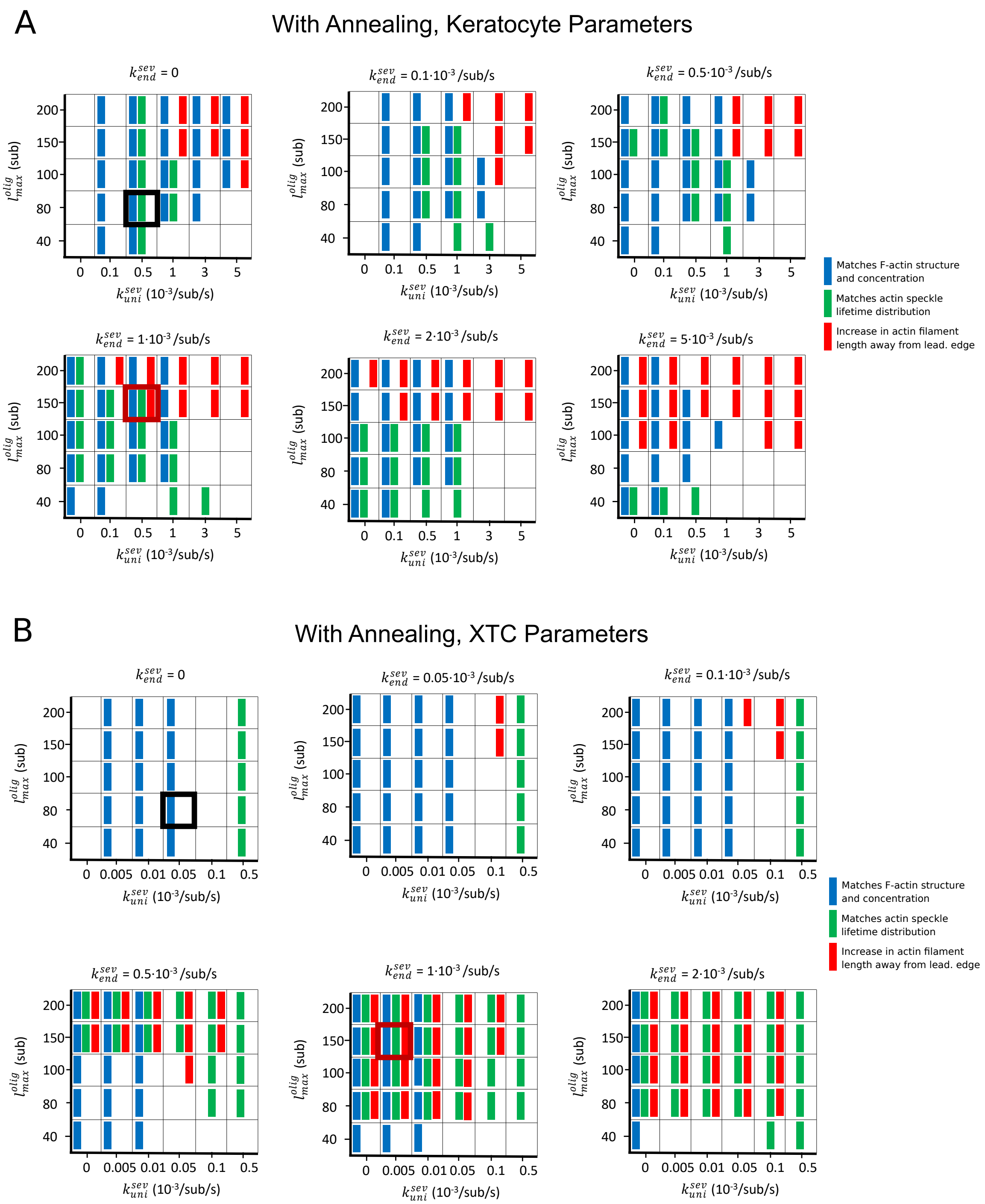


Figure S5: Parameter scan of uniform and enhanced end severing rates ($k_{unif}^{sev}$, $k_{end}^{sev}$) and maximum oligomer length ($l_{max}^{olig}$), with oligomer annealing but no debranching $k_{debr} = 0$. Other parameters are listed in Table 1. Boxes highlighted with thick black and red frames correspond to the parameters used in the black and red curves of Figs. 4 and 5. Blue rectangles indicate that F-actin, barbed end, and branch concentration profiles agree with experimental observations. Green rectangles indicate that speckle lifetimes, appearance and disappearance locations agreed with the results of SiMS measurements. Red rectangles indicate an increase in length between the filament lengths measured in $0 - 1\mu m$ and $3 - 4\mu m$. (A) Keratocyte parameter scan. (B) XTC cell parameter scan.

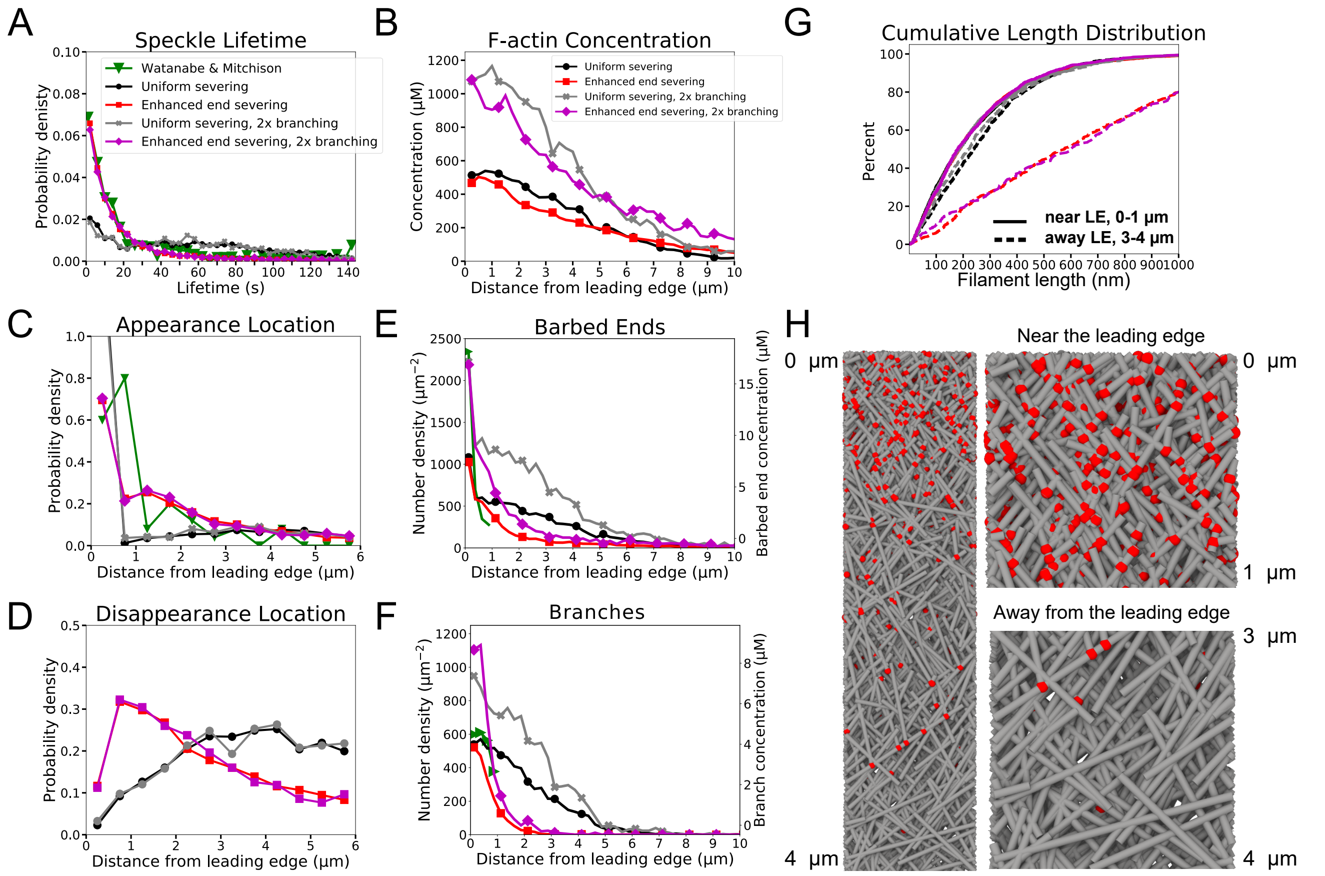


Figure S6: (A-G) Repeat of simulations of Fig. 5 (XTC cells) with doubled branching rate $k_{br}$. Doubling the branching rate in the XTC parameter set results in concentrations closer to prior estimates for XTC cells but the branch distribution deviates from the measurements of Vinzenz et al. for fibroblasts. Black and red curves are the same data as in Fig. 5 for uniform or enhanced end severing, respectively. Gray curves show the case of uniform severing as in the black curve, but with doubled $k_{br}$. Magenta curves show the case of enhanced severing as in the red curve but with doubled $k_{br}$. Gray and magenta results are from one simulation evolved to steady state. (H) Simulation snapshot for case of enhanced end severing and doubled $k_{br}$ as in magenta curves.

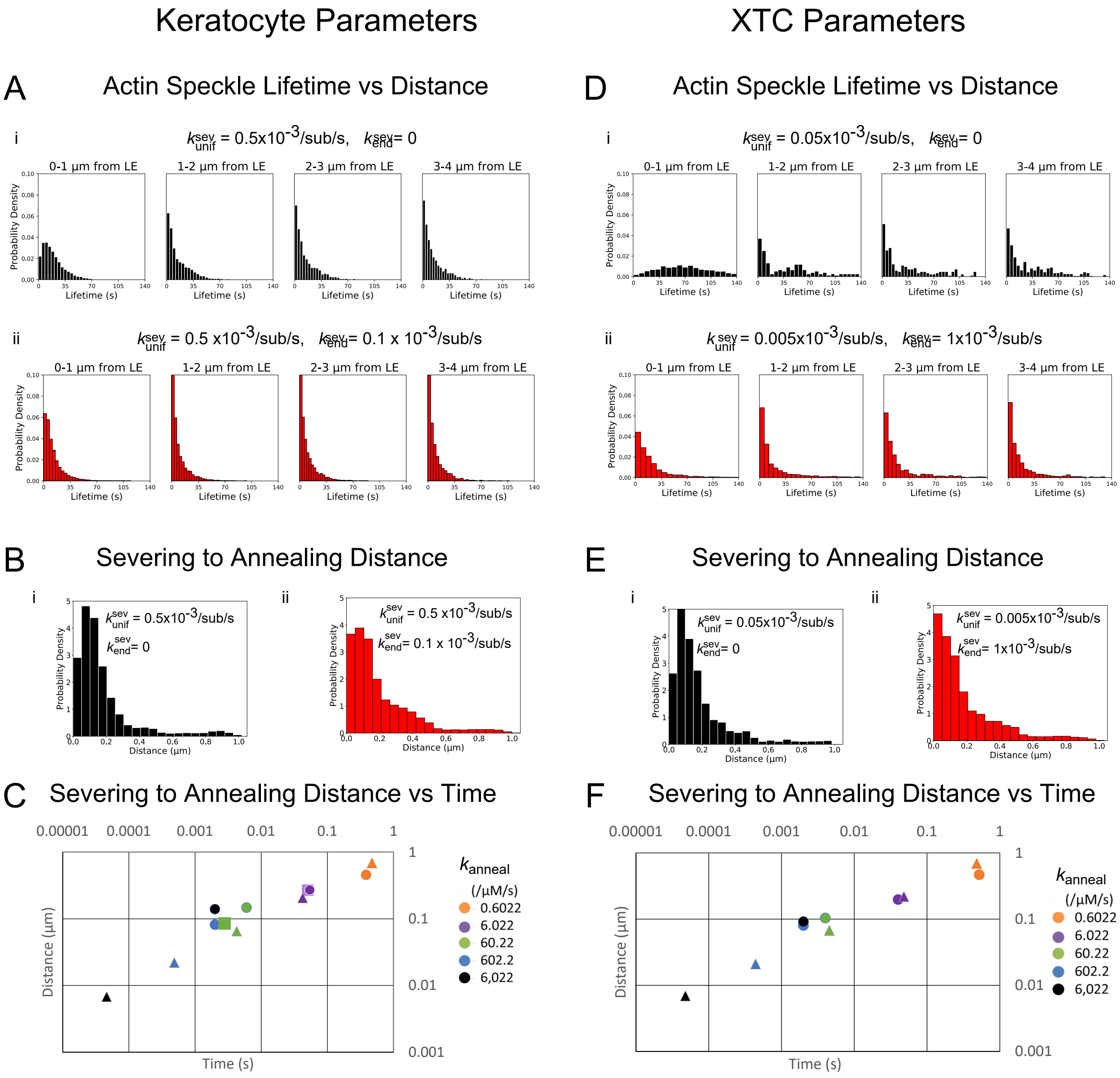


Figure S7: Quantification of actin turnover in simulations with severing and annealing of Fig. 4 (keratocyte parameters A-C) and Fig. 5 (XTC parameters, D-F) from 5 simulations in steady-state over 20 s each. (A and D) Probability density of simulated actin speckle lifetimes separated into 1 $\mu$m regions based on their appearance location for parameters with annealing and uniform severing (i, black) or with enhanced end severing (ii, red). (B and E) Probability density of the distance between the location the oligomer was created (within 4 $\mu$m of the leading edge) and annealing position for parameters with annealing and uniform severing (i, black) or with enhanced end severing (ii, red). (C and F) Log-log plot of the median distance traveled by an oligomer in between severing and annealing versus the median time spent diffusing, for various annealing rate constants. This plot serves as a check of the effect smallness of simulation time step. Triangles are the expected values from $r = (4D_{olig}t)^{1/2}$ with $t = k_{anneal}C_B{}^{-1}$ and $C_B$ is average concentration of free barbed ends. Circles are values from simulations with the standard time step, $2 \cdot 10^{-3}$ s. Squares show simulations with a smaller time step of $2 \cdot 10^{-4}$ s. The simulated lamellipodium width was 1 $\mu m$. For $k_{anneal} > 602.2$ $\mu$M$^{-1}$s$^{-1}$, the simulation results are limited by the chosen time step. The smaller annealing rate constant $k_{anneal} = 6.02$ $\mu$M$^{-1}$s$^{-1}$ has better agreement with the theoretical expectation. (F) Used a maximum oligomer length $l_{max}^{olig} = 40$ sub.
